# A new bioinformatic analysis highlights a subpopulation of mitochondria with highly methylated DNA present in oocytes, early embryos and blood

**DOI:** 10.64898/2026.10.26.761975

**Authors:** Camila Bruna de Lima, Mohamed Oudihat, Marcella Pecora Milazzotto, Marc-André Sirard

**Author notes:** **Corresponding author**: Marc-André Sirard Département des Sciences Animales, Université Laval, Québec, QC, Canada.

## Abstract

**Background:** Mitochondria are present in virtually all eukaryotic cells, yet several aspects of their function remain poorly understood. Although their activity depends on more than 1,000 genes, the mitochondrial genome encodes only 38 genes, including 13 protein-coding genes. Mitochondrial DNA (mtDNA) differs substantially from nuclear DNA (nDNA), notably because it lacks histones and contains virtually no intronic or intergenic sequences. DNA methylation of mtDNA has been reported in oocytes, embryos, and various human diseases, occurring not only at CpG sites but also in other cytosine contexts. However, average methylation levels are generally considered to be very low. By re-examining methylation patterns at the individual-read level, this study highlights marked heterogeneity in mtDNA methylation and proposes a potential explanation for previously conflicting observations.

**Results:** Using bovine oocytes and embryos, as well as blood samples, we analyzed multiple datasets generated by whole-genome EM-seq, which captures both nuclear and mitochondrial DNA methylation. Sequenced reads were separated and analyzed individually to quantify methylation levels either at specific cytosines or across all cytosines within each read. The resulting distributions consistently displayed a U-shaped pattern, where the vast majority of reads were completely unmethylated while a small subset was highly methylated. The proportion of methylated reads increased from oocytes to 8-16 cell embryos, then declined in blastocysts and blood samples. This distribution pattern was highly conserved among samples from the same developmental stage and remained remarkably similar across embryonic stages, while differing from blood. When only heavily methylated reads (>50% methylation) were analyzed, their methylation profiles along the 16 kb mitochondrial genome were even more conserved across tissues. These observations suggest the existence of distinct mtDNA populations with different methylation states that are largely obscured by conventional bioinformatic approaches based on averaged methylation levels.

**Conclusions:** The identification of highly methylated mtDNA reads reveals heterogeneity among mtDNA molecules and raises the hypothesis that these molecules may be associated with functionally distinct mitochondrial states. The heavily methylated population, termed “mitoQueens,” may exhibit reduced gene expression, lower metabolic activity, decreased reactive oxygen species (ROS) production, and potentially reduced mutation accumulation. In contrast, the predominantly unmethylated population, referred to as “mitoWorkers,” would be responsible for most energy production and cellular functions. Similar to stem cells, “Queen” mitochondria may represent a reserve population that can be activated under specific conditions and preserved for transmission to future generations.

**Summary sentence:** Oocytes, embryos, and mammalian blood contain a subpopulation of mitochondria with elevated DNA methylation, which may remain inactive and serve as a pristine template for later mitochondrial replication.

## Introduction

Mitochondria play a central role in oocyte growth, maturation, and early embryonic development, providing ATP, buffering calcium, and regulating redox balance. Unlike somatic cells, oocyte mitochondria display unique structural and functional characteristics that reflect the dual requirement for energy production and long-term genetic preservation. Accumulating evidence indicates that oocytes harbor distinct mitochondrial populations that differ in localization, ultrastructure, and metabolic activity [1]. Early electron microscopy studies in mammalian oocytes revealed heterogeneous mitochondrial morphologies, with small, round mitochondria dispersed throughout the cytoplasm and larger mitochondrial aggregates localized near the nucleus or cortex [2]. These dispersed mitochondria are characterized by poorly developed cristae and low electron density, features associated with reduced oxidative phosphorylation capacity. Such mitochondria are considered metabolically quiescent, a state thought to limit reactive oxygen species (ROS) production and protect mitochondrial DNA from damage [3]. In contrast, a second population forms spatially organized clusters, frequently associated with the endoplasmic reticulum, generating specialized mitochondria–ER complexes. These complexes have been shown to function as calcium signaling hubs, coordinating Ca²⁺ release during fertilization and oocyte activation [4]. Live-cell imaging studies using mitochondrial membrane potential-sensitive dyes demonstrated that clustered mitochondria exhibit higher ΔΨm than dispersed mitochondria, supporting functional specialization [5]. It is also known that the small, mostly rounded oocyte mitochondria contain only one or 2 copies of mtDNA compared to somatic cells [6].

In oocytes, ATP imaging and fluorescent biosensors further revealed that ATP-rich microdomains coincide with mitochondrial aggregates, particularly in regions supporting spindle formation and chromosome segregation [7]. This metabolic polarization suggests that localized mitochondrial populations selectively support energy-demanding processes, such as meiotic progression and fertilization. Molecular studies provide additional support for mitochondrial heterogeneity. Differences in mitochondrial DNA replication timing and transcriptional activity have been observed between mitochondrial subpopulations during oocyte growth, indicating functional compartmentalization at the genomic level [8,9]. In bovine, porcine, and human oocytes, progressive mitochondrial clustering correlates with oocyte developmental competence, linking specific mitochondrial populations to embryo viability [10].

Collectively, these findings support a model in which oocytes contain a reserve population of quiescent mitochondria that safeguards mitochondrial inheritance, alongside a strategically localized, highly active population that drives calcium signaling, ATP production, and early developmental events. This dual-mitochondrial system may represent a fundamental adaptation of the mammalian oocyte, ensuring both immediate functional performance and long-term developmental fidelity.

Recent evidence has revealed that mitochondrial DNA (mtDNA) represents an additional layer of epigenetic regulation during bovine preimplantation development. Genome-wide analyses have demonstrated that mtDNA is methylated across the mitochondrial genome and that these methylation patterns undergo dynamic remodeling from the oocyte through the blastocyst stage [11]. Importantly, mitochondrial methylation is not randomly distributed but shows gene-specific variations, suggesting regulated epigenetic control of mitochondrial function during early development [11]. Furthermore, mtDNA methylation in bovine oocytes and embryos is strand-biased, predominantly affecting the heavy strand, indicating a structured and potentially functional epigenetic organization of the mitochondrial genome [11]. In parallel, embryos exhibiting different developmental kinetics display distinct DNA methylation signatures associated with mitochondrial and metabolic pathways, linking mitochondrial epigenetic regulation to developmental competence [12]. Together, these findings establish that mitochondrial methylation is developmentally regulated, sensitive to early embryonic dynamics, and likely involved in controlling mitochondrial activity during early development. This emerging concept of mitoepigenetic regulation prompts a reassessment of mitochondrial contributions to embryo quality and highlights mtDNA methylation as a potential mediator of early developmental programming.

One unresolved feature of mitochondrial DNA methylation is the very low proportion of methylated cytosines relative to nuclear genomic sequences, together with the presence of non-CpG methylation, which has been observed in mouse and bovine oocytes [12,11]. In our previous studies [11], we also observed an atypical distribution of methylation, in which most reads showed nearly 0% methylation, whereas a distinct subpopulation of reads displayed methylation levels above 25%. This observation prompted us to investigate whether two populations of mitochondrial DNA molecules exist, each with a distinct methylation signature and possibly a different fate or function. Using MethPat, an alternative bioinformatic approach to explore read-level methylation patterns, we detected this dichotomy in EM-seq data from both oocytes and somatic lineages. These analyses are supporting a hypothesis of the existence of “mitoqueens” with a function differing from “mitoWorkers” to ensure the transmission of minimal mutation of the mitochondrial genome across generations. Figure 1A illustrates the 2 types of mitochondrial DNA that would exist in the same cell.

**Figure 1:**
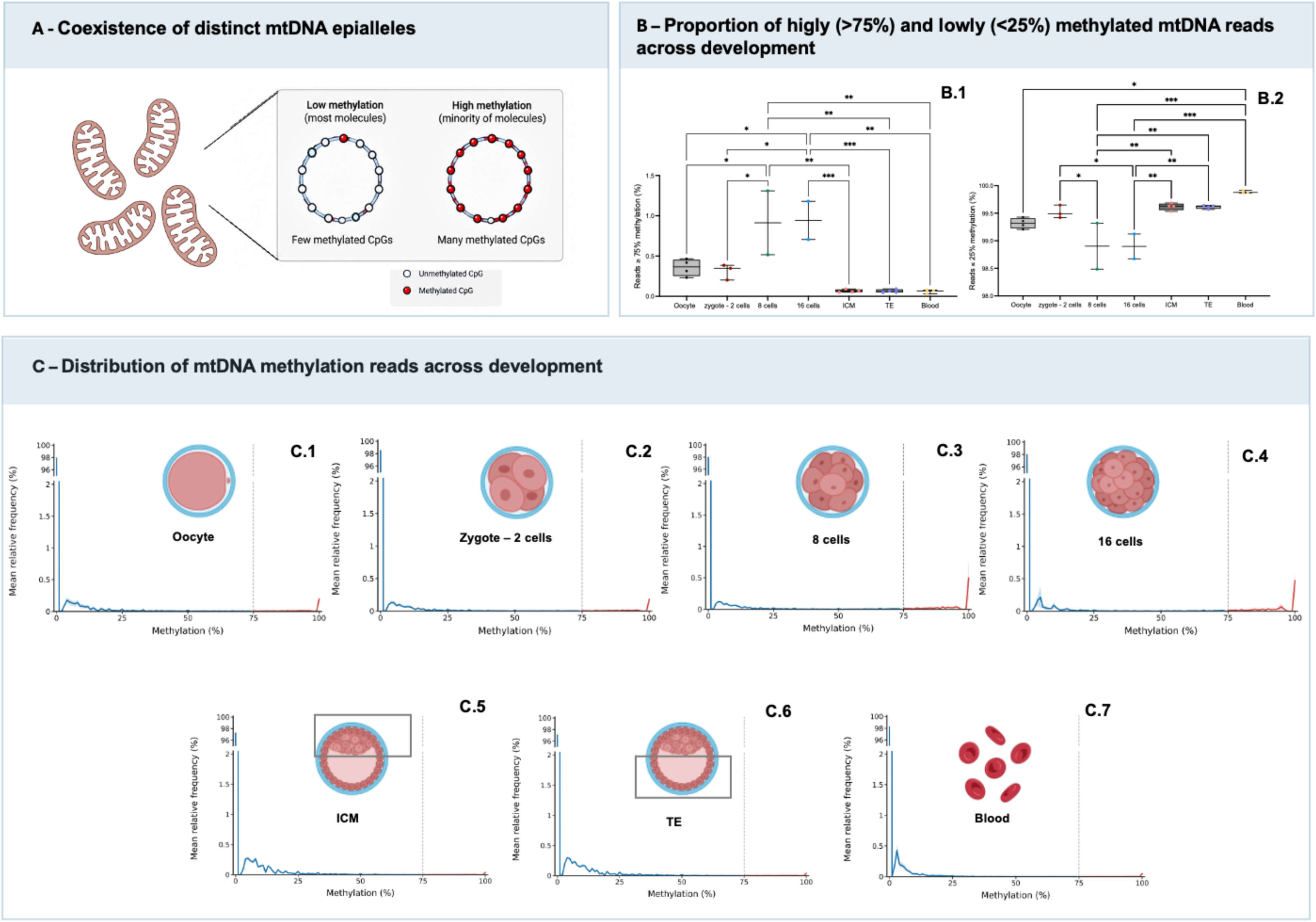
The distribution of mitochondrial DNA (mtDNA) methylation levels was analyzed at the individual read level for each group. Reads were classified according to their methylation percentage, ranging from 0% to 100%, and grouped into two categories: non-Queen reads (<25% methylation) and Queen reads (≥75% methylation). For each biological replicate, the relative frequency of reads at each methylation level was calculated as the number of reads within that category divided by the total number of analyzed reads, multiplied by 100. The mean relative frequency across the two biological replicates was plotted, with shaded areas representing the standard deviation (SD). The dashed vertical line indicates the 75% methylation threshold. A discontinuous y-axis was used to improve visualization of low-frequency methylation categories. No inferential statistical tests were performed. Statistical analysis indicates differences (* for p<0.05, ** P<0.01 and *** for p< 0.001). Panel C represent the distribution of methylated reads on the mitochondrial genome from oocytes to blastocysts and blood.

## Materials and Methods

The analyzed samples were generated using EM-seq sequencing. They included Bos taurus oocytes, 2-cell and 8-cell embryos, 16-cell embryos, blastocysts as inner cell mass (ICM) or trophectoderm (TE) and DNA extracted from blood cells.

### Oocyte and embryo production and collection

Ovaries were obtained from a commercial slaughterhouse and transported to the laboratory for follicular aspiration using a syringe and needle. Cumulus–oocyte complexes (COCs) were selected and subjected to *in vitro* maturation and fertilization according to previously established protocols [11]. A subset of oocytes was mechanically denuded by repeated pipetting and collected for subsequent analyses. After fertilization, presumptive zygotes were denuded by vortex and cultured in 90 µL droplets of ECS50-oviduct medium supplemented with 3% BSA and covered with mineral oil at 38.5 °C, 6% CO₂, 5% O₂, and high humidity. On Day 4, embryos were transferred to ECS50-uterus medium supplemented with 3% BSA and cultured until Day 7. Embryos at the 2-, 8-, and 16-cell stages, as well as expanded blastocysts, were collected according to their developmental stage [11]. For blastocyst lineage separation, the trophectoderm (TE) was first mechanically removed, followed by purification of the inner cell mass (ICM) by immunosurgery. Briefly, expanded blastocysts were treated with pronase (10 mg/mL) to remove the zona pellucida and incubated for 1 h at 38.5 °C in 20% anti-bovine serum diluted in SOF-HEPES. After washing, embryos were incubated for 1 h at 38.5 °C with 20% guinea pig complement in SOF-HEPES. The remaining cells were repeatedly pipetted to remove residual TE cells and obtain purified ICMs. Collected oocytes, cleavage-stage embryos, TE, and purified ICM samples were stored at −80 °C until subsequent molecular analyses [11].

### DNA extraction and EM-seq library preparation

Total DNA was extracted from pools of five oocytes or embryos and from whole blood across four biological replicates using the Quick-DNA Microprep Kit (Zymo Research), following proteinase K digestion. Libraries were prepared using the NEBNext® Enzymatic Methyl-seq Kit (New England Biolabs) according to the manufacturer’s instructions. Lambda DNA and pUC19 were included as unmethylated and methylated controls, respectively, to assess conversion efficiency. DNA was fragmented to approximately 300 bp, enzymatically converted, indexed, and amplified. Library quality and concentration were assessed using an Agilent TapeStation (D1000HS), and sequencing was performed on an Illumina NovaSeq 6000 platform using paired-end 100-bp reads [11].

### Methylation extraction and calling for EM-seq data

Raw sequencing reads were first processed using fastp v0.24.0 [17] to remove adapter sequences and low-quality bases and to perform quality filtering. The mitochondrial reference genome (16,338 bp; NC_006853.1) and the complete *Bos taurus* reference genome (ARS-UCD2.0, July 1, 2023) were prepared using the Bismark genome preparation procedure, generating the C-to-T and G-to-A converted reference sequences. To prevent the inclusion of reads originating from nuclear mitochondrial DNA segments (NUMTs) [18), the trimmed reads were first aligned to the mitochondrial reference genome (16,338 bp; NC_006853.1) using Bismark v0.24.1 [13). The resulting aligned reads were then converted from BAM to FASTQ format using *SAMtools* [15] and realigned to the complete *Bos taurus* reference genome (ARS-UCD2.0, July 1, 2023). Only reads that are aligned to the mitochondrial chromosome in this second alignment were retained for downstream analysis. Following alignment, duplicate reads were removed using the Bismark deduplication procedure to generate BAM files containing uniquely retained, non-duplicated reads for downstream analyses. Methylation calling was subsequently performed using the Bismark methylation extractor. This generated read methylation extraction files containing read-level methylation information and CX reports providing methylation information at single-cytosine resolution, including genomic position, strand, methylated reads, unmethylated reads, and sequence context (CpG, CHG, or CHH). These files were subsequently used for downstream characterization and quantification of mitochondrial DNA methylation.

### Cytosine- and Read-Level DNA Methylation in Mitochondrial DNA Samples

In this section, two types of mtDNA methylation data were analyzed. First, cytosine-level methylation was assessed at each cytosine position and represented as the percentage of methylated and unmethylated reads aligned to a given cytosine position represented in the CX reports. Read-level DNA methylation was calculated from the methylation extraction files generated by Bismark, which contained methylation information derived from reads aligned to the C-to-T and G-to-A converted reference sequences. A custom extraction pipeline was developed to calculate the average DNA methylation at the read level for each sample. Read methylation levels were calculated by determining methylation-state information encoded in the XM tag of each aligned read in methylation extraction files for each sample. In these files, methylated CpG sites are encoded as Z and unmethylated CpG sites as z; methylated CHG sites are encoded as X and unmethylated CHG sites as x; and methylated CHH sites are encoded as H and unmethylated CHH sites as h [13]. For each read, the methylation level was calculated as the proportion of methylated cytosines among all informative cytosines within that read, separately for CpG, CHG, and CHH contexts and for all cytosine contexts combined. The resulting read-level methylation values were then summarized across reads to obtain mean methylation levels for each context.

### Reads and Individual Cytosine-Level Methylation Profiling of Mitochondrial DNA Samples

MethPat [14] was used to summarize DNA methylation patterns derived from the Bismark methylation extractor output. For each amplicon, the genomic positions of methylated cytosines, the observed DNA methylation patterns, and their corresponding abundance were extracted and summarized. The resulting information was compiled into a tab-delimited text file and subsequently used for downstream analyses and visualization of read-level DNA methylation patterns across EM-seq samples. In the resulting visualizations, each column represents an observed methylation pattern, with a histogram below the plot showing the number of reads displaying the same pattern. Each cell represents the methylation state at a CpG position within an individual read: methylated CpGs are shown in yellow, unmethylated CpGs in blue, and other bases corresponding to non-cytosine positions in gray. Each row represents a CpG position along the mitochondrial genome. The proportion column indicates the fraction of reads at each cytosine position classified as methylated, unmethylated, or unknown.

The R package **Circlize** (v0.4.18; [16]) was used to generate circular plots of hypermethylated reads, defined as reads with a mean methylation level >50%, for each analyzed sample and across the three cytosine contexts (CpG, CHG, and CHH). The circular plots were used to visualize read alignments across the mitochondrial genome and their distribution relative to mitochondrial gene annotations. The genomic positions of each mitochondrial gene were obtained from the NCBI annotation of the *Bos taurus* mitochondrial reference genome (16,338 bp; NC_006853.1). In addition, Circlize was used to visualize cytosine-level methylation percentages, their genomic positions across the mitochondrial genome, and their distribution relative to mitochondrial gene annotations. For this purpose, a specific annotation file containing the genomic position and methylation information for each cytosine was generated.

### Comparative Analysis of Mitochondrial DNA Methylation Profiles in all stages using high vs low methylated reads

The proportion of highly and lowly methylated mitochondrial DNA (mtDNA) reads was determined for each biological replicate as the percentage of reads exhibiting ≥75% or <25% methylation, respectively, relative to the total number of analyzed mtDNA reads. Differences among developmental stages and tissues (oocytes, zygote to 2 cell embryos, 8-cell embryos, 16-cell embryos, inner cell mass [ICM], trophectoderm [TE], and blood) were evaluated using one-way analysis of variance (ANOVA), followed by Tukey’s post hoc test for multiple comparisons. Statistical significance was set at p < 0.05. Data are presented as boxplots showing the median and interquartile range, with individual biological replicates represented by colored dots. To assess whether differences in sequencing depth could influence the observed proportions of methylated reads, a random down sampling analysis was performed using 2,000 reads per biological replicate, repeated 100 times independently. The proportion of methylated reads was recalculated for each iteration to evaluate the robustness of the observed patterns. The down sampling analysis yielded results consistent with those obtained from the complete datasets, indicating that the observed differences were not driven by variations in sequencing depth (data not shown).

### Comparative Analysis of Mitochondrial DNA Methylation Profiles using all reads

Comparative analyses of mitochondrial DNA (mtDNA) methylation profiles were performed in R between blood and oocyte samples, as well as between the inner cell mass (ICM) and trophectoderm (TE). Mean cytosine methylation levels were calculated for each sample across the three cytosine sequence contexts (CpG, CHG, and CHH). Methylation levels were visualized using boxplots generated with the *ggplot2* R package. Statistical differences between groups were assessed using the Wilcoxon rank-sum test, with significance determined based on the resulting *P*-values.

## Results

### Cytosine methylation level in EMseq-analysed Oocytes

The distribution of mtDNA methylation levels was analyzed at single-cytosine resolution across all cytosine contexts (CpG, CHG, and CHH) in the mitochondrial genome of oocytes. The data were generated using EM-seq. For each cytosine, the methylation ratio was calculated as the number of methylated reads divided by the total number of methylated and unmethylated reads. The methylation distribution was strongly skewed toward unmethylated reads (Figure 1, C-1-6), indicating that most cytosines analyzed were either unmethylated or exhibited very low methylation levels. This pattern was consistently observed across all available samples. The proportion of highly methylated reads ( >75%) goes up at the 8-cells and 16-cells embryos to around 1 % and decrease in later stages and in blood. In all stages, the distributions appeared relatively bimodal, with a U-shaped pattern beginning to emerge at approximately 0.2 (or 20%). This difference may be attributable, at least in part, to the greater abundance of mitochondrial reads in oocytes and in 8-cell and 16-cell embryos compared with ICM, TE, and blood samples (suppl Table 1).

### Mean cytosine methylation level in EM-seq oocytes

The distribution of DNA methylation at the single-cytosine level across all cytosine contexts (CpG, CHG, and CHH) revealed highly consistent methylation patterns across the different samples analyzed. Notably, similar methylation patterns were observed in several mitochondrial regions, including the D-loop 1, tRNA-Leu, ND4, and sRNA regions. These observations suggest that methylation patterns established in the oocyte may be maintained during early embryonic development and persist across subsequent developmental stages (Figure 2).

**Figure 2.**
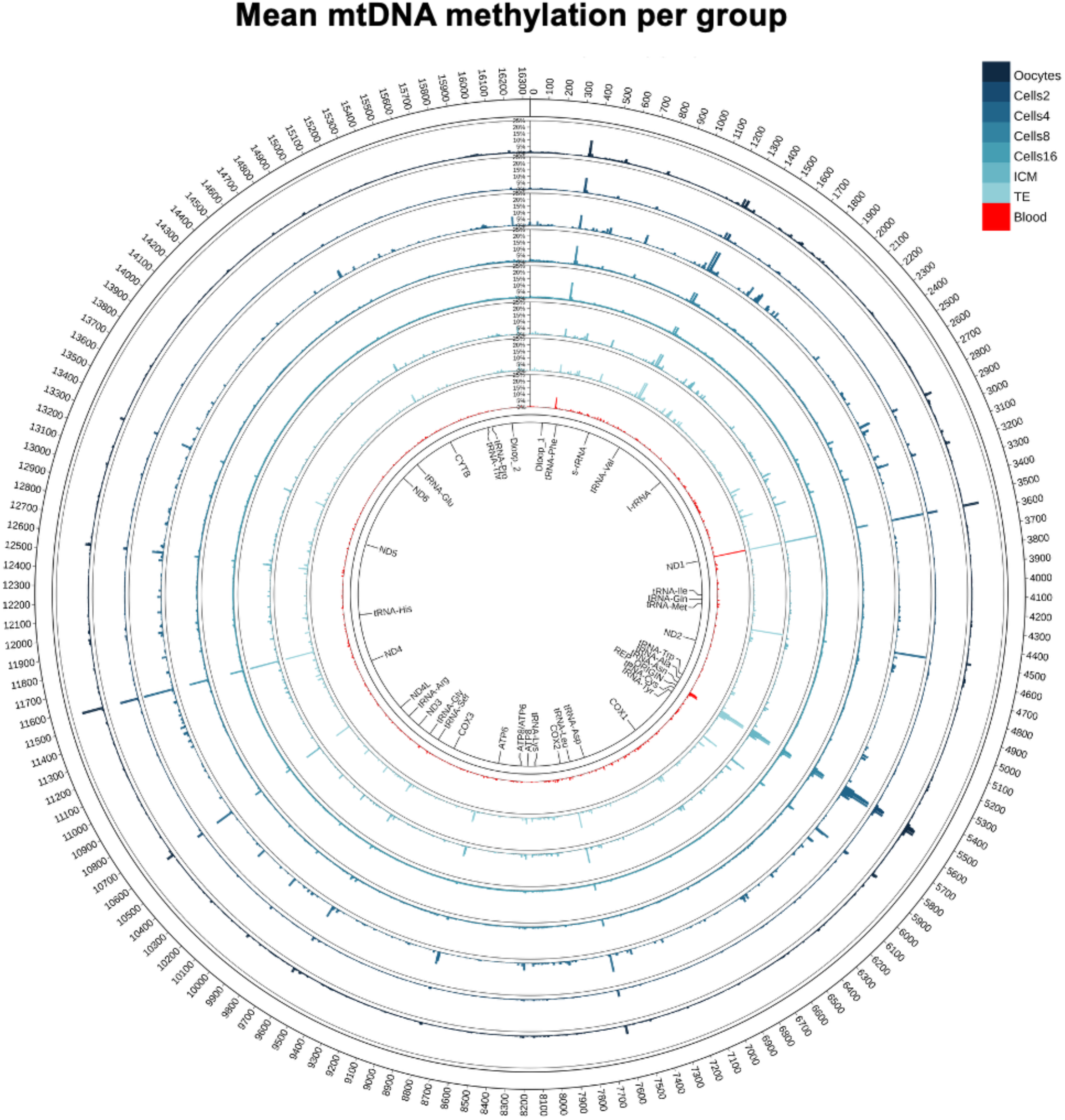
Mitochondrial cytosine methylation levels across genomic positions and developmental stages. Methylation profiles across mitochondrial cytosine positions show consistent patterns across the different embryonic developmental stages, with similar positional methylation patterns observed between samples. A similar methylation pattern was also observed in blood, suggesting that several positional features of mitochondrial cytosine methylation are maintained across developmental stages and tissues. Blood in red and oocytes to embryos in blue.

### Identification of highly methylated reads in EMseq oocyte samples

Most studies investigating mtDNA methylation with conventional sequencing approaches report methylation as the average percentage of methylated and unmethylated reads at individual cytosine sites. However, this approach does not capture positional information encoded within epiallelic methylation patterns. To overcome this limitation, we used MethPat to visualize methylation patterns at the epiallelic level. As shown in Figure 3, this analysis revealed distinct methylation patterns within the ATP6 and COX1 region in oocytes, zygote to 2 cells, 8-cells, 16 cells embryos, and blastocysts. Reads with 100% CpG methylation are shown in red, whereas reads with unmethylated CpG sites are shown in blue. We interpret the highly methylated COX1 epiallele as a candidate Mitoqueen-associated mtDNA molecule, while recognizing that this designation remains operational and requires independent validation by long-read sequencing, mitochondrial enrichment, and functional assays.

**Figure 3.**
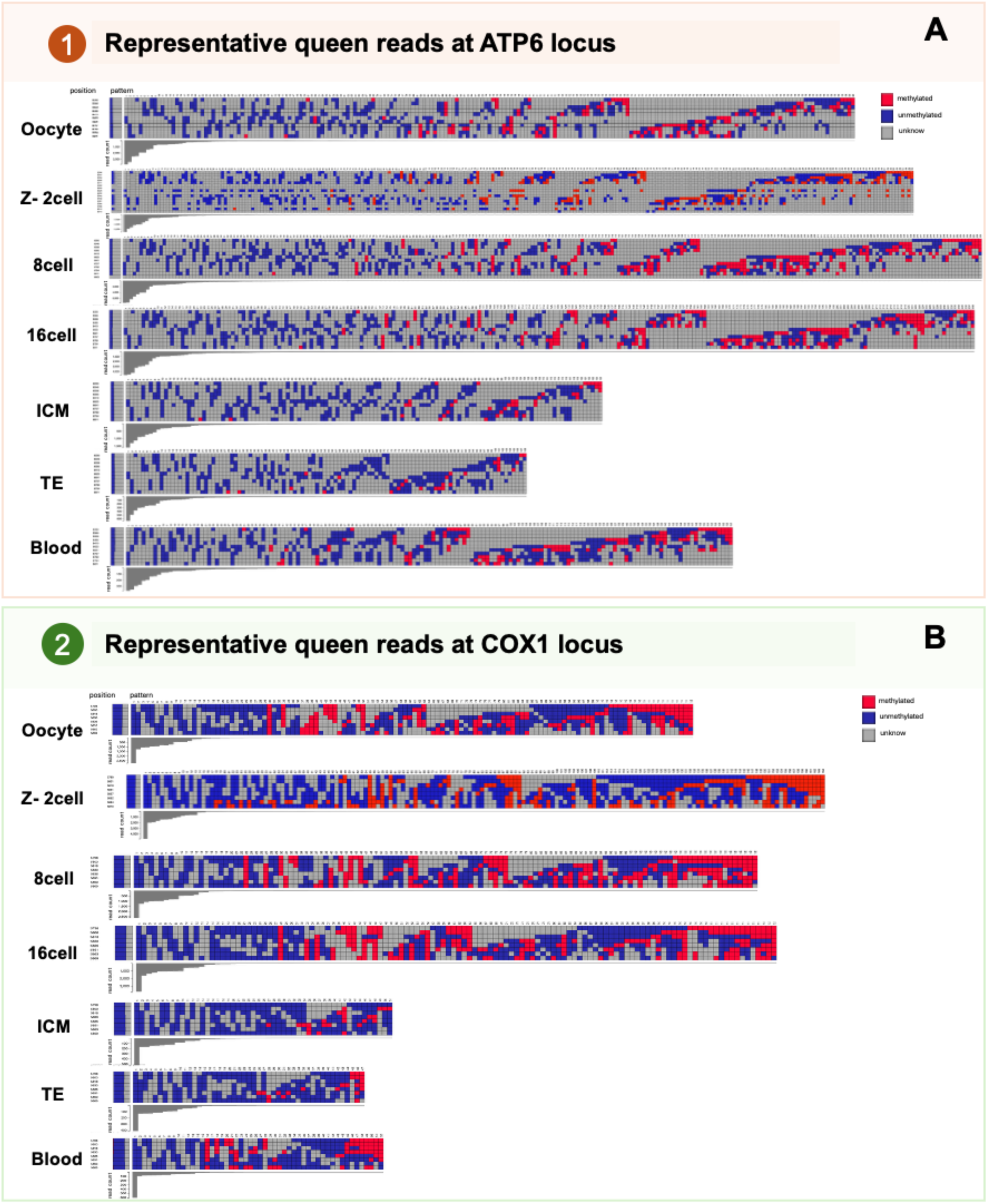
Read-level mitochondrial DNA methylation patterns across the ATP6 and COX1 loci during bovine embryonic development and in blood samples. Representative epiallelic methylation patterns were visualized using MethPat based on enzymatic methyl-sequencing (EM-seq) data from different developmental stages and sample types, including oocytes, 2-cell, 8-cell, and 16-cell embryos, the inner cell mass (ICM), trophectoderm (TE), and blood samples. **(A)** Methylation patterns within the mitochondrial ATP6 locus. **(B)** Methylation patterns within the mitochondrial COX1 locus. Each horizontal row represents an aggregated CpG methylation pattern, while columns correspond to successive cytosine positions within the analyzed region. Methylated cytosines are represented in red, unmethylated cytosines in blue, and positions with undetermined methylation status in gray. The horizontal arrangement of colored positions illustrates the co-occurrence and spatial distribution of methylated and unmethylated cytosines within the COX1 and ATP6 loci. The gray histograms displayed beneath each methylation matrix represent the number of sequencing reads (*read counts*) associated with each methylation pattern, with taller bars indicating more frequently observed patterns. Additionally, the first box at the beginning of each row indicates the proportions of methylated and unmethylated CpG sites within the corresponding pattern, as well as positions for which methylation status could not be determined, particularly when a nucleotide other than cytosine was observed, represented in gray. Across the developmental stages examined, methylated epiallelic patterns were identified in both mitochondrial loci, some of which may represent candidate Mitoqueen-associated mitochondrial DNA molecules.

**Figure 4.**
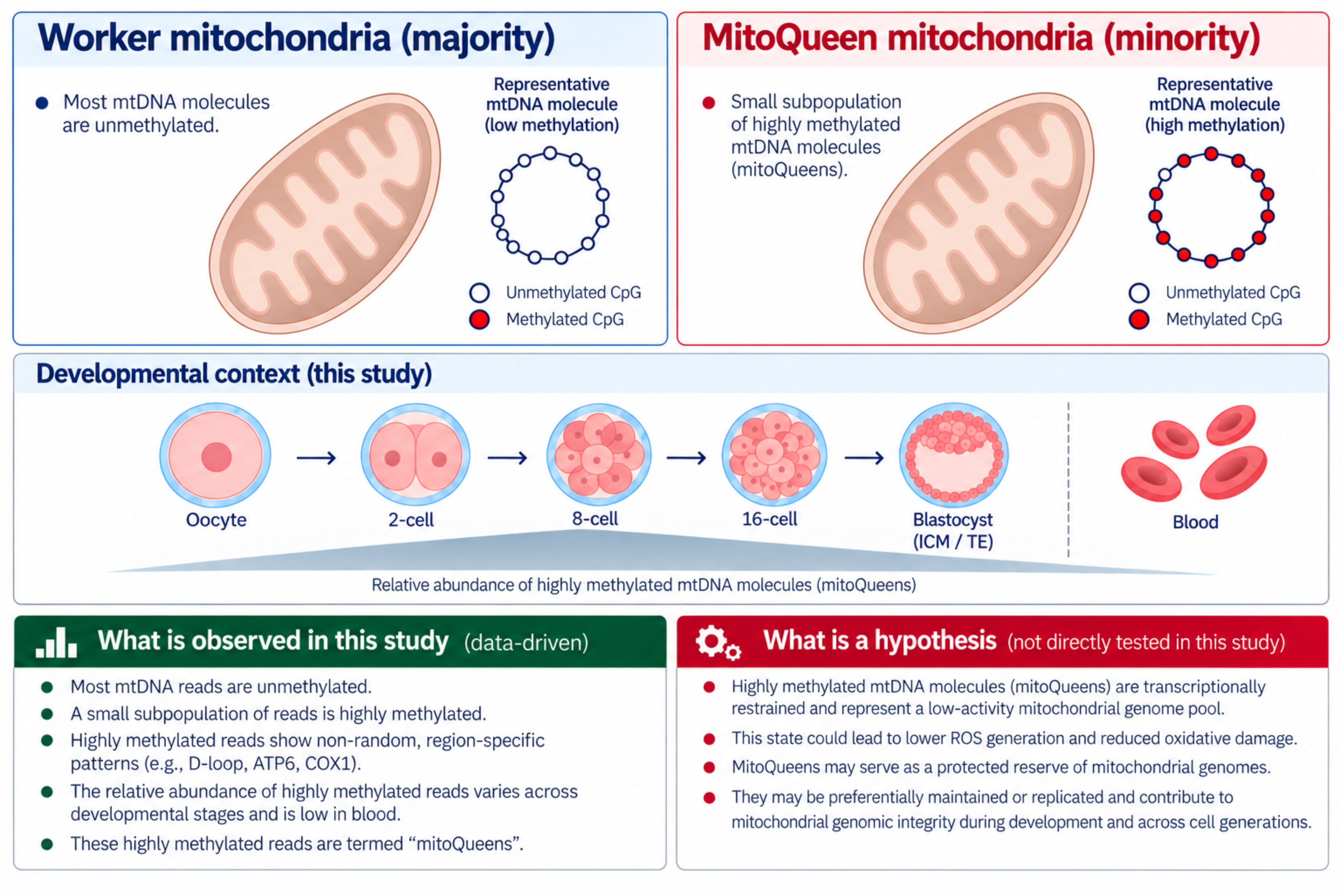
Graphical abstract.

To identify Mitoqueen’s presence exhibiting a hypermethylated profile, we further investigated the presence of highly methylated reads that could potentially correspond to Mitoqueen copies (see methods). The results presented in supplementary table 1 were derived from read-level methylation metrics calculated across all analyzed EMeq samples. Among the samples examined, the oocytes and embryos at the 8-cell and 16-cell stages displayed a relatively high number of reads (more than 15,000) with methylation levels exceeding 50%.

This observation may be explained, at least, by the greater number of reads aligned to the mitochondrial genome in oocytes and early embryos at the 8-cell and 16-cell stages (suppl table 1). Thus, the higher number of highly methylated reads observed in oocytes and early embryos may reflect their greater mitochondrial sequencing depth, which could in turn be associated with a higher mitochondrial genome copy number in oocytes and during the early stages of embryonic development. We assessed if the distribution of methylation was different between ICM (inner cell mass) and TE (Trophectoderm) and inner cell mass cells contain more methylated sequences (supplementary figure 1).

Reads with more than 50% methylation were then selected and visualized in circos plots to determine whether methylation followed a distinctive distribution across the mitochondrial genome (supplementary figure 2). To illustrate the maintenance of a similar pattern across the developmental stages, an example of each stage is provided to compare with the blood sample (supplementary figure 3). Notably, methylation was enriched in several mitochondrial regions, including D-loop 1, tRNA-Leu, ND4, and sRNA regions, consistent with the positional patterns shown in Figure 2 but restricted here to reads with more than 50% methylation.

### Oocyte vs Blood

To assess whether methylation levels differ between oocytes and somatic tissues, we used blood as a readily available source of mtDNA and compared the mean of three blood samples with three oocyte samples. supplementary figure 3 illustrates that methylation level in average is lower in blood than in oocytes.

## Discussion

Conventional bioinformatic approaches that average methylation levels across cytosine positions can obscure biologically important read-level variation. In this study, we provide evidence that mitochondrial DNA methylation is not evenly distributed across mtDNA copies but is enriched in a small subset of highly methylated reads while most reads remain unmethylated. This interpretation is supported by the convergence of short-read EM-seq data, read-level pattern analysis and NUMT-aware filtering [18]. Although further functional validation will be required, these results suggest that average mtDNA methylation values may mask biologically relevant subpopulations of mitochondrial genomes.

Our results also suggest some dynamic regulation of the highly methylated portions of the mtDNA with a rise between the oocyte and genome activation (8-16 cell stage in bovine) and a reduction as the cells differentiate in TE and eventually in blood. These remaining methylated sequenced are therefore potentially conserve in all cells, although a confirmation from each tissue would be required. In fact, the literature already reports low level of methylation in most tissues tested for different human diseases (see below) but it seems that no one has reorganized the data to look at specific reads instead of the mean which mask most of it.

### The controversy about mtDNA methylation

The existence and functional relevance of mitochondrial DNA (mtDNA) methylation remain highly debated within the epigenetics field. Early studies argued that mtDNA is essentially unmethylated, citing the absence of canonical CpG islands and histone-based chromatin structure as evidence against classical epigenetic regulation [19,20]. In addition, technical limitations of bisulfite sequencing, incomplete conversion, alignment ambiguity, and nuclear mitochondrial DNA sequence contamination have raised concerns that reported mtDNA methylation signals could be artefactual [19,20,21]. These concerns are directly relevant to the present study because the highly methylated molecules represent only a small fraction of the total mtDNA reads. For this reason, the present analysis applied stringent NUMT filtering, evaluated conversion controls, normalized sequencing depth across samples, and analyzed methylation at the individual-read level [18].

In contrast, a growing number of studies using improved methodological controls have reported low but reproducible levels of mtDNA methylation, often in a strand-biased and region-specific manner [11,24,25]. Genome-wide analyses further demonstrated that mtDNA methylation patterns are developmentally regulated and responsive to environmental conditions such as in vitro culture, arguing against random technical noise [11]. The identification of mitochondrial-localized DNMTs and TET enzymes has also provided mechanistic plausibility for active mitochondrial epigenetic regulation [24,25,26]. Nevertheless, critics argue that mtDNA methylation levels are often orders of magnitude lower than nuclear DNA methylation, questioning whether such small differences can exert meaningful biological effects [19,21,22,23]. Supporters counter that mitochondrial biology is highly sensitive to threshold effects and copy-number amplification, allowing subtle epigenetic changes to influence mitochondrial transcription and cellular metabolism disproportionately, as observed for mutant mtDNA copies [27]. As a result, the debate has shifted from whether mtDNA methylation exists to whether and how it is functionally interpreted by the cell.

### Human diseases

Although disease-associated differences in mitochondrial DNA (mtDNA) methylation are often quantitatively small, frequently below 1–5%, several mechanisms may explain why such changes could still be biologically relevant. First, mitochondria exist in hundreds to thousands of copies per cell, meaning that a small average shift in mtDNA methylation could reflect coordinated changes across many mitochondrial genomes or, alternatively, larger changes within a small subpopulation of mtDNA molecules [8,27]. Either scenario could affect mitochondrial transcriptional output if the affected molecules are functionally specialized or preferentially replicated [8,11]. Because mtDNA encodes essential subunits of the oxidative phosphorylation system, even modest transcriptional effects may influence ATP production, redox balance, and metabolic signaling [28]. Also, mtDNA methylation is non-random and region-specific, as also shown in this study. Genome-wide mitochondrial methylome analyses have shown that methylation differences are concentrated at specific genes and exhibit strand bias, particularly on the heavy strand, suggesting targeted regulation rather than stochastic noise [11,12]. Thus, small average differences may mask larger functional effects at critical regulatory loci.

Together, these references support the idea that modest mtDNA methylation differences can be biologically meaningful and relevant to disease risk. An additional explanation is that only a small subset of mitochondria may harbor these methylation changes, potentially affecting replication control or the function of a protected mitochondrial genome. The results presented here raise an alternative possibility: low average mtDNA methylation may reflect a small subpopulation of highly methylated mitochondrial genomes and not low-level methylation uniformly distributed across the mitochondrial population.

### The potential role of methylation on mtDNA

The functional significance of the highly methylated mtDNA molecules remains unknown. Mitochondrial DNA methylation has been associated with reduced transcription of mitochondrial-encoded genes, particularly when methylation occurs near promoter-like regions on the heavy strand, which contains the majority of protein-coding genes [11,25,22]. Experimental and correlative studies indicate an inverse relationship between mtDNA methylation and mitochondrial transcript abundance, supporting a potential repressive role of methylation on mtDNA transcription [11,25,22].

Among the few mitochondrial genes, those encoding complex I (NADH: ubiquinone oxidoreductase) subunits appear particularly sensitive. Reduced expression of complex I genes has been linked to increased mtDNA methylation, leading to a limitation in electron entry into the respiratory chain [11,24,25]. Because complex I activity is a rate-limiting step for oxidative phosphorylation, even modest transcriptional repression can significantly lower ATP production at the cellular level [28].

Reduced electron flux through the respiratory chain also leads to lower mitochondrial membrane potential and decreased thermogenic output, particularly in tissues where mitochondrial uncoupling contributes to heat generation [20,28]. In parallel, decreased electron transport reduces electron leakage from the respiratory chain, resulting in lower reactive oxygen species (ROS) generation, especially at complex I, a primary site of mitochondrial ROS production [29]. Importantly, reduced ROS generation has protective consequences for the mitochondrial genome. ROS are a major source of oxidative damage to mtDNA, which lacks histone protection and has limited repair capacity compared with nuclear DNA [29,30]. By dampening respiratory chain activity, mtDNA methylation may indirectly reduce mutation rates and preserve mitochondrial genomic integrity over time [27,29,30].

Taken together, these observations support a model in which highly methylated mtDNA molecules could represent a transcriptionally restrained mitochondrial genome population. However, the present study does not establish whether these molecules are less transcriptionally active, experience lower oxidative damage, or preferentially persist or replicate. Functional experiments will therefore be required to determine whether methylation is a driver, marker, or consequence of such a protected state.

mtDNA methylation is associated with a low-activity mitochondrial state characterized by reduced transcription, diminished complex I expression, lower ATP and heat production, decreased ROS generation, and potentially reduced mutational burden. At present, the data supports an association between methylation patterns and mitochondrial regulatory states rather than proving direct causality. In this framework, mitochondrial methylation may act as a protective epigenetic tuning mechanism that balances bioenergetic output with long-term mitochondrial genome stability, but functional perturbation experiments will be required to determine whether methylation is a driver, a marker, or a consequence of this state.

Several limitations should be considered when interpreting these findings. First, the frequency of highly methylated reads is low, making the analysis sensitive to sequencing depth, conversion efficiency, base-calling accuracy, and alignment stringency. Second, although NUMT-aware filtering reduces the likelihood of nuclear contamination, it cannot eliminate all mapping ambiguity. Finally, the proposed Mitoqueen model remains a working hypothesis that will require orthogonal validation, including targeted long-read sequencing, mitochondrial enrichment, single-cell or single-organelle approaches, and functional assays linking methylation state to transcription, replication, oxidative stress, and developmental outcome.

### Our hypothesis

Although the results of this paper require more validation and extension to nanopore sequencing, they suggest a new hypothesis implying that oocytes may contain a subpopulation of mitochondria carrying more highly methylated mtDNA molecules that could contribute to the preservation of mitochondrial genomic integrity, consistent with literature on mitochondrial heterogeneity, oocyte mitochondrial activity, and the mitochondrial genetic bottleneck [1,8,9,31]. In somatic cells, a comparable small subpopulation of methylated mtDNA molecules could serve as a protected reserve that is preferentially maintained or replicated during cell division [1,8,31]. In this model, most mitochondria remain unmethylated and metabolically active, whereas a minority exist in a lower-activity, methylated state, the Queens.

By maintaining a pool of epigenetically restrained mitochondria, the cell could minimize oxidative damage to mtDNA, which lacks histone protection and has limited repair capacity [29,30]. This mechanism would reduce mutation accumulation, particularly during replicative expansion, and preserve mitochondrial genomic integrity across cell generations. Such a strategy aligns with threshold models of mitochondrial biology, where small changes in mitochondrial activity have disproportionate effects on cellular stress and long-term viability [27,28].

In a recent paper on human embryos [32) the authors have characterized the gene expression originating from the mitochondrial genome in single cell essays and concluded that the distribution of mitochondria was different in cells which become trophectoderm versus the inner cell mass cells [32). The difference was associate with more oxidative phosphorylation permissive genes in the trophoblast cells. This is additional indirect evidence that the methylation controlling gene expression is potentially differently distributed between cells lines. Overall, this hypothesis frames mtDNA methylation as a selective, stress-induced tuning mechanism, enabling somatic cells to balance metabolic output with mitochondrial genome protection, especially during division and environmental challenge.

## Acknowledgements

The research results presented here were supported by NSERC in Canada and FAPESP in Brazil. The initial sequencing from blood was supported by Genome Quebec and Genome Canada in a different project.

## Data Availability

Data related to this study are available under GEO accession number **GSE2G0131 (blood) oocytes and embryos (GSE230476)**

---

**Suppl fig 1 Comparison of distribution mtDNA of methylated reads between ICM (inner cell mass) and TE (trophectoderm) cells samples.** The figure shows a statistically significant difference in methylation distribution, with higher mean DNA methylation levels observed in ICM compared with TE samples (p<0.0001)

**Suppl fig 2. Distribution of highly methylated reads across without NUMT mitochondrial genome in EM-seq samples from different Oocytes.** The figure shows the distribution of reads with high methylation levels (> 50%) across the mitochondrial genome in EM-seq samples from Oocytes to blood. The outer circle represents mitochondrial genomic positions. Blue points indicate read methylation levels in the CpG context, orange points represent read methylation levels in the CHG context, and red points represent read methylation levels in the CHH context. The central region indicates mitochondrial gene coordinates.

**Suppl fig 3. Comparison of mean DNA methylation profiles between blood and oocyte samples.** The figure shows a statistically significant difference in methylation profiles, with higher mean DNA methylation levels observed in oocytes compared with blood samples.

**Suppl Table 1:** Distribution of samples data, coverage, methylation means and number of highly methylated reads.

## Notes

**Grant support:** The authors would like to thank the Natural Sciences and Engineering Research Council of Canada (NSERC) for supporting this work and the São Paulo Research Foundation (FAPESP) # 2023/15734-1 and # 2021/11747-6.

### Competing Interest Statement

The authors have declared no competing interest.

## References

1. Bahety D, Böke E, Rodríguez-Nuevo A. Mitochondrial morphology, distribution and activity during oocyte development. Trends Endocrinol Metab. 2024;35(10):902–917. doi: 10.1016/j.tem.2024.03.002. PMID: 38599901.

2. Hyttel P, Callesen H, Greve T. Ultrastructural features of preovulatory oocyte maturation in superovulated cattle. J Reprod Fertil. 1986;76(2):645–656. doi: 10.1530/jrf.0.0760645. PMID: 3084771.

3. Jones A, Van Blerkom J, Davis P, Toledo AA. Cryopreservation of metaphase II human oocytes effects mitochondrial membrane potential: implications for developmental competence. Hum Reprod. 2004;19(8):1861–1866. doi: 10.1093/humrep/deh313. PMID: 15229209.

4. Dumollard R, Marangos P, Fitzharris G, Swann K, Duchen M, Carroll J. Sperm-triggered [Ca2+] oscillations and Ca2+ homeostasis in the mouse egg have an absolute requirement for mitochondrial ATP production. Development. 2004;131(13):3057–3067. doi: 10.1242/dev.01181. PMID: 15163630.

5. Wilding M, Dale B, Marino M, di Matteo L, Alviggi C, Pisaturo ML, Lombardi L, De Placido G. Mitochondrial aggregation patterns and activity in human oocytes and preimplantation embryos. Hum Reprod. 2001;16(5):909–917. doi: 10.1093/humrep/16.5.909. PMID: 11331637.

6. Shoubridge EA, Wai T. Mitochondrial DNA and the mammalian oocyte. Curr Top Dev Biol. 2007;77:87–111. doi: 10.1016/S0070-2153(06)77004-1. PMID: 17222701

7. Yu Y, Dumollard R, Rossbach A, Lai FA, Swann K. Redistribution of mitochondria leads to bursts of ATP production during spontaneous mouse oocyte maturation. J Cell Physiol. 2010;224(3):672–680. doi: 10.1002/jcp.22171.

8. Wai T, Teoli D, Shoubridge EA. The mitochondrial DNA genetic bottleneck results from replication of a subpopulation of genomes. Nat Genet. 2008;40(12):1484–1488. doi: 10.1038/ng.258. PMID: 19029901.

9. Woods DC, Khrapko K, Tilly JL. Influence of maternal aging on mitochondrial heterogeneity, inheritance, and function in oocytes and preimplantation embryos. Genes (Basel). 2018;9(5):265. doi: 10.3390/genes9050265. PMID: 29883421; PMCID: PMC5977205.

10. Stojkovic M, Machado SA, Stojkovic P, Zakhartchenko V, Hutzler P, Gonçalves PB, Wolf E. Mitochondrial distribution and adenosine triphosphate content of bovine oocytes before and after in vitro maturation: correlation with morphological criteria and developmental capacity after in vitro fertilization and culture. Biol Reprod. 2001;64(3):904–909. doi: 10.1095/biolreprod64.3.904.

11. de Lima CB, Martin H, Pecora Milazzotto M, Sirard MA. Genome-wide methylation profile of mitochondrial DNA across bovine preimplantation development. Epigenetics. 2023;18(1):2241010. doi: 10.1080/15592294.2023.2241010.

12. Shirane K, Toh H, Kobayashi H, Miura F, Chiba H, Ito T, Kono T, Sasaki H. Mouse oocyte methylomes at base resolution reveal genome-wide accumulation of non-CpG methylation and role of DNA methyltransferases. PLoS Genet. 2013;9(4):e1003439. doi: 10.1371/journal.pgen.1003439. PMID: 23637617; PMCID: PMC3630097.

13. Krueger F, Andrews SR. Bismark: a flexible aligner and methylation caller for Bisulfite-Seq applications. Bioinformatics. 2011;27(11):1571–1572. doi: 10.1093/bioinformatics/btr167. PMID: 21493656; PMCID: PMC3102221.

14. Wong NC, Pope BJ, Candiloro IL, Korbie D, Trau M, Wong SQ, et al. MethPat: a tool for the analysis and visualisation of complex methylation patterns obtained by massively parallel sequencing. BMC Bioinformatics. 2016;17:98. doi: 10.1186/s12859-016-0950-8.

15. Petr Danecek, James K Bonfield, Jennifer Liddle, John Marshall, Valeriu Ohan, Martin O Pollard, Andrew Whitwham, Thomas Keane, Shane A McCarthy, Robert M Davies, Heng Li, Twelve years of SAMtools and BCFtools, GigaScience, Volume 10, Issue 2, February 2021, giab008, 10.1093/gigascience/giab008.

16. Gu Z, Gu L, Eils R, Schlesner M, Brors B. circlize Implements and enhances circular visualization in R. Bioinformatics. 2014 Oct;30(19):2811–2. doi: 10.1093/bioinformatics/btu393. Epub 2014 Jun 14. PMID: 24930139

17. Chen S, Zhou Y, Chen Y, Gu J. fastp: an ultra-fast all-in-one FASTQ preprocessor. Bioinformatics. 2018;34(17):i884–i890. doi: 10.1093/bioinformatics/bty560.

18. Cox SN, Varvara AS, Pesole G. MitSorter: a standalone tool for accurate discrimination of mtDNA and NuMT ONT reads based on differential methylation. Bioinformatics Advances. 2025;5(1):vbaf135. doi: 10.1093/bioadv/vbaf135. PMID: 40688360; PMCID: PMC12275464.

19. Mechta M, Ingerslev LR, Fabre O, Picard M, Barrès R. Evidence suggesting absence of mitochondrial DNA methylation. Front Genet. 2017;8:166. doi: 10.3389/fgene.2017.00166. PMID: 29163634; PMCID: PMC5671948.

20. Yin J, Gong Z, Mao S. Apparent methylation of mitochondrial DNA in WGBS is dominated by technical artifacts. Epigenetics Chromatin. 2026. doi: 10.1186/s13072-026-00692-z.

21. Basrai S, Bahcheli AT, Tan D, Zuzarte PC, Bevan A, Chan T, et al. Multi-platform reassessment of human mitochondrial DNA methylation reveals signals consistent with technical artifacts. bioRxiv. 2026. doi: 10.64898/2026.06.10.730935. Preprint.

22. Yin MT, Guo L. Mitochondrial DNA methylation: state-of-the-art in molecular mechanisms and disease implications. J Adv Res. 2026;83:455–473. doi: 10.1016/j.jare.2025.08.029. PMID: 40848899; PMCID: PMC13131407.

23. Picard M, Wallace DC, Burelle Y. The rise of mitochondria in medicine. Mitochondrion. 2016;30:105–116. doi: 10.1016/j.mito.2016.07.003.

24. Quiros PM, Mottis A, Auwerx J. Mitonuclear communication in homeostasis and stress. Nat Rev Mol Cell Biol. 2016;17:213–226. doi: 10.1038/nrm.2016.23.

25. Shock LS, Thakkar PV, Robinson JM, Taylor SM. Cytosine methylase and hydroxymethylase activity in mammalian mitochondria. Front Cell Dev Biol. 2025;13:1677402. doi: 10.3389/fcell.2025.1677402.

26. Saini SK, Mangalhara KC, Prakasam G, Bamezai RNK. DNA methyltransferase 1 isoform 3 methylates mitochondrial genome and modulates its biology. Sci Rep. 2017;7(1):1525. doi: 10.1038/s41598-017-01743-y. PMID: 28484249; PMCID: PMC5431478.

27. Wallace DC, Chalkia D. Mitochondrial DNA genetics and the heteroplasmy conundrum in evolution and disease. Cold Spring Harb Perspect Biol. 2013;5(11):a021220. doi: 10.1101/cshperspect.a021220.

28. Spinelli JB, Haigis MC. The multifaceted contributions of mitochondria to cellular metabolism. Nat Cell Biol. 2018;20:745–754. doi: 10.1038/s41556-018-0124-1.

29. Murphy MP. How mitochondria produce reactive oxygen species. Biochem J. 2009;417(1):1–13. doi: 10.1042/BJ20081386. PMID: 19061483; PMCID: PMC2605959.

30. Shokolenko IN, Wilson GL, Alexeyev MF. Aging: a mitochondrial DNA perspective, critical analysis and an update. World J Exp Med. 2014;4(4):46–57. doi: 10.5493/wjem.v4.i4.46. PMID: 25414817; PMCID: PMC4237642.

31. Cao L, Shitara H, Sugimoto M, Hayashi JI, Abe K, Yonekawa H. New evidence confirms that the mitochondrial bottleneck is generated without reduction of mitochondrial DNA content in early primordial germ cells of mice. PLoS Genet. 2009;5(12):e1000756. doi: 10.1371/journal.pgen.1000756. PMID: 19997484.

32. Piasecki AJ, Franco M, Capelluto F, Khrapko K, Tilly JL, Woods DC.Mitochondria-associated transcription precedes oxidative phosphorylation in human embryos. Reproduction. 2026 Oct 5;172(4):xaag117. doi:10.1093/reprod/xaag117. PMID: 42752920.

