## Supplementary figures and images for "A new bioinformatic analysis highlights a subpopulation of mitochondria with highly methylated DNA present in oocytes, early embryos and blood"

### Suppl fig 1

Suppl Fig 1

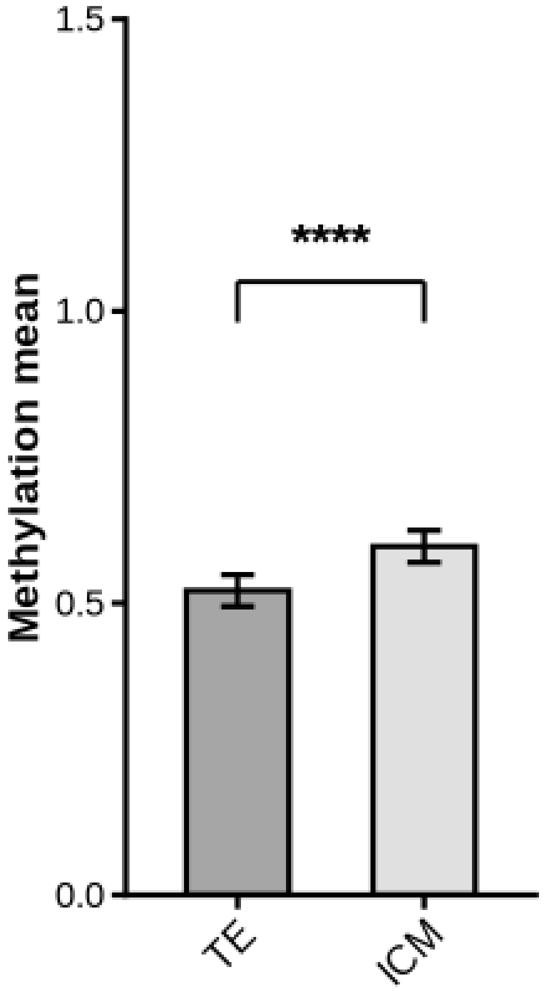

### Suppl fig 3

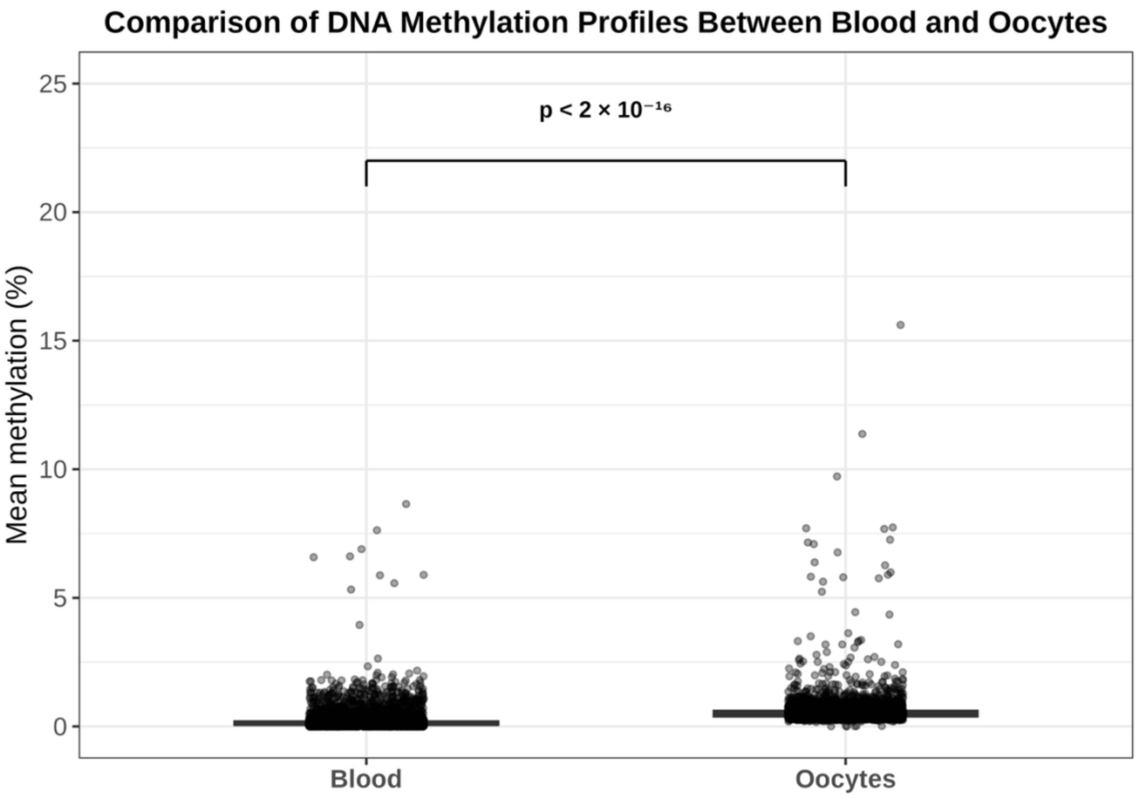
