## Supplementary material for "A new bioinformatic analysis highlights a subpopulation of mitochondria with highly methylated DNA present in oocytes, early embryos and blood": Suppl fig 2

Suppl figure 2

Examples of methylation patterns using reads with > 50% methylation

Oocyte

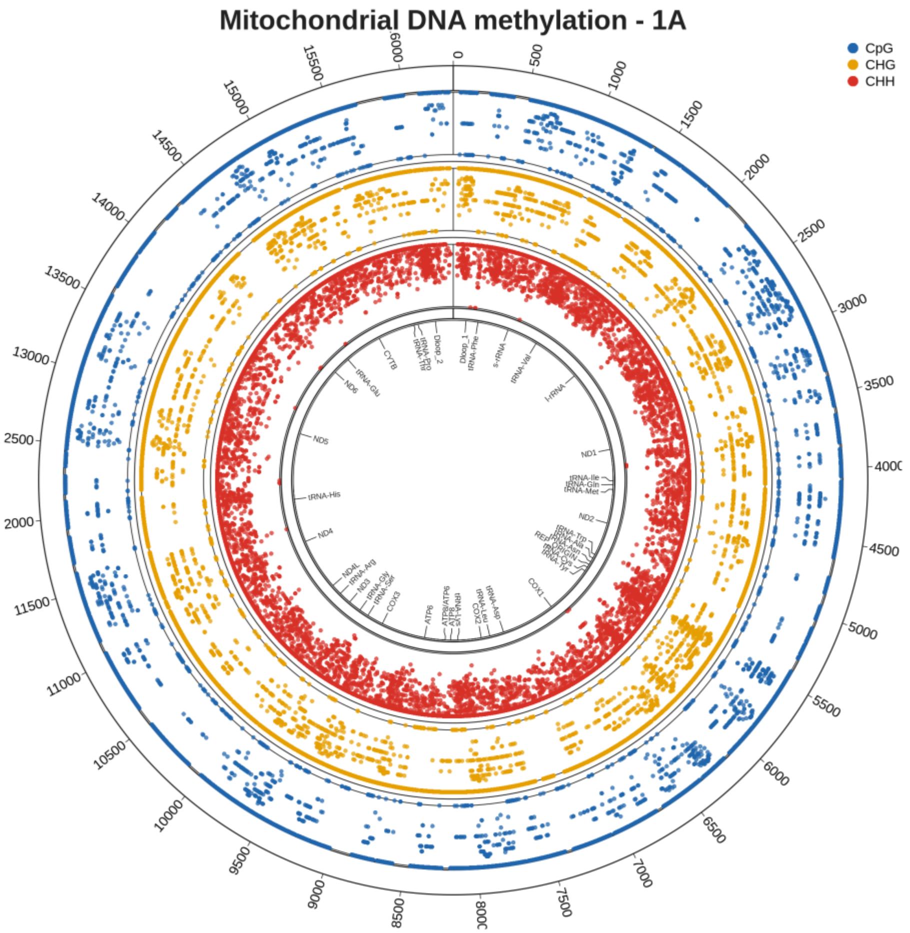


2 cells

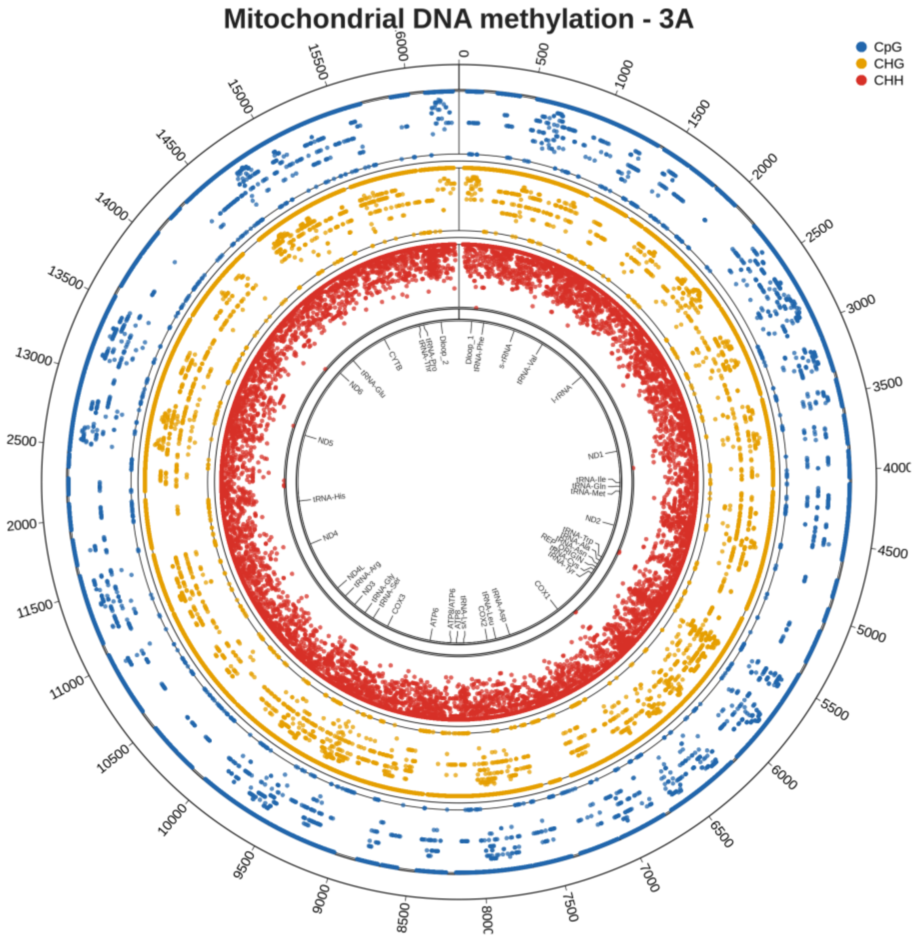

8 cells

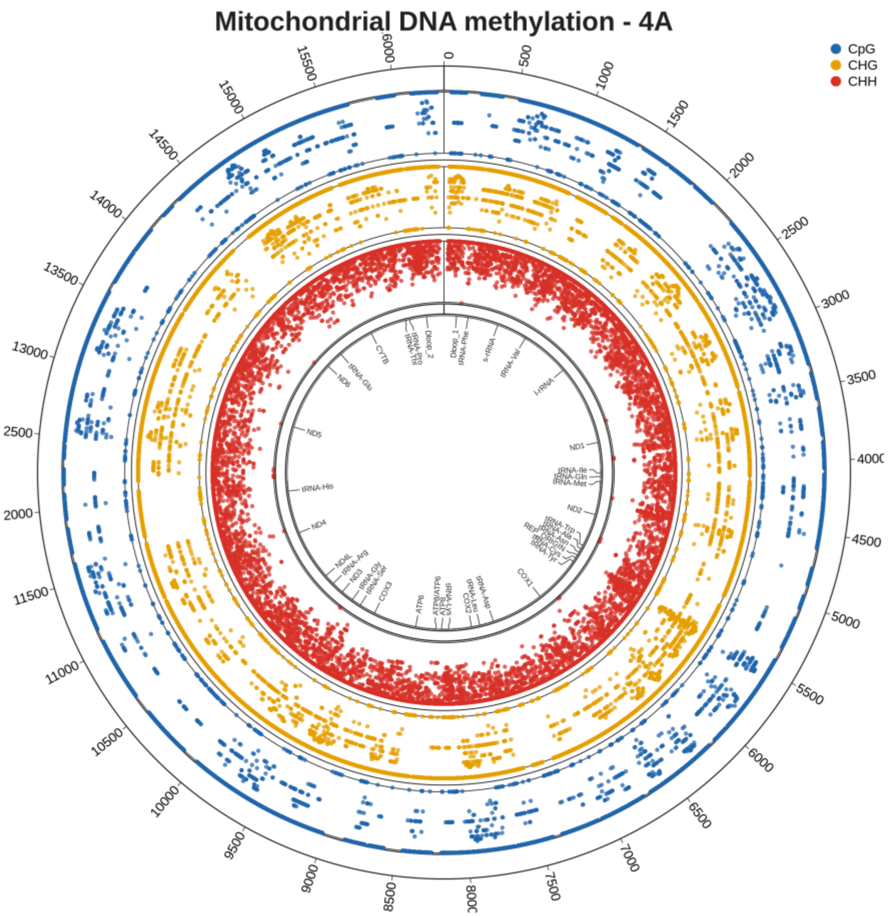


16 cells

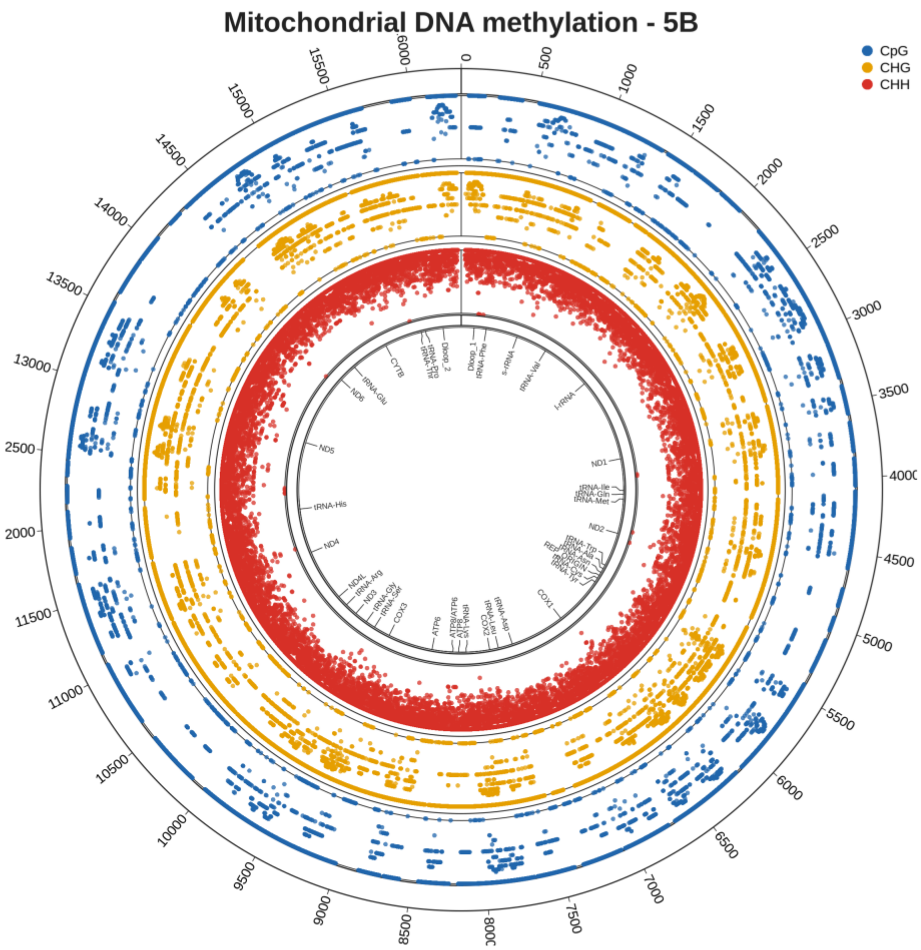

ICM

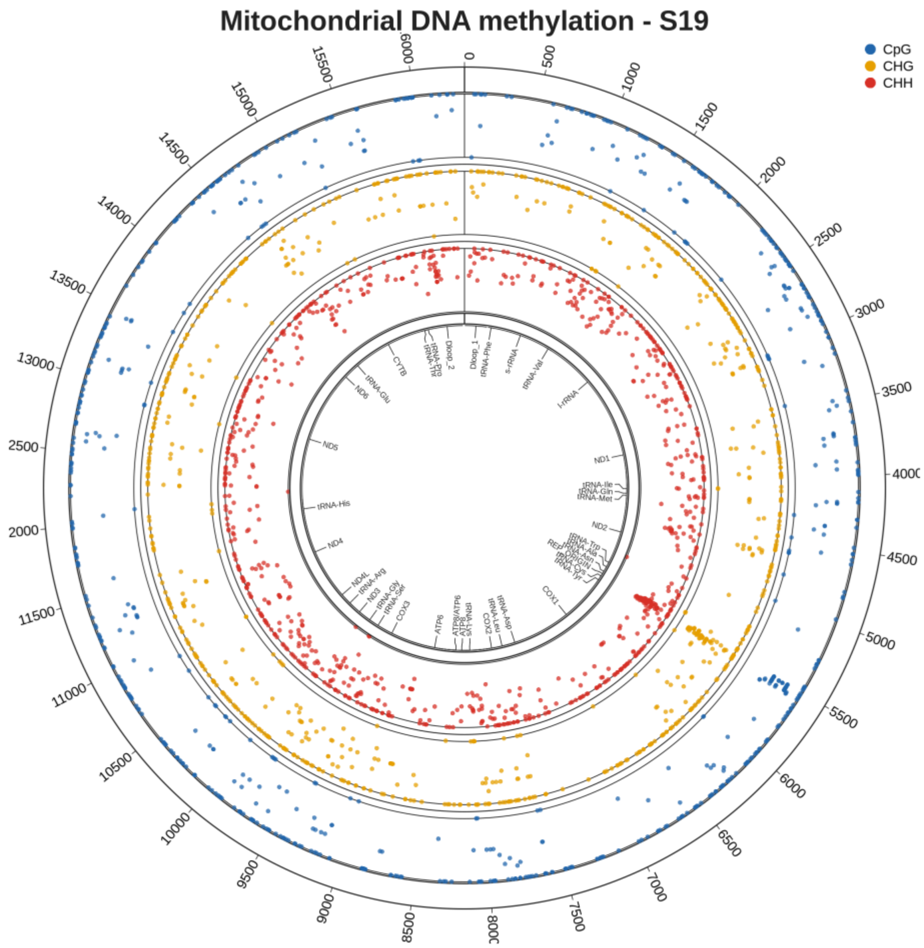
Mitochondrial DNA methylation - S19
CpG
CHG
CHH

TE

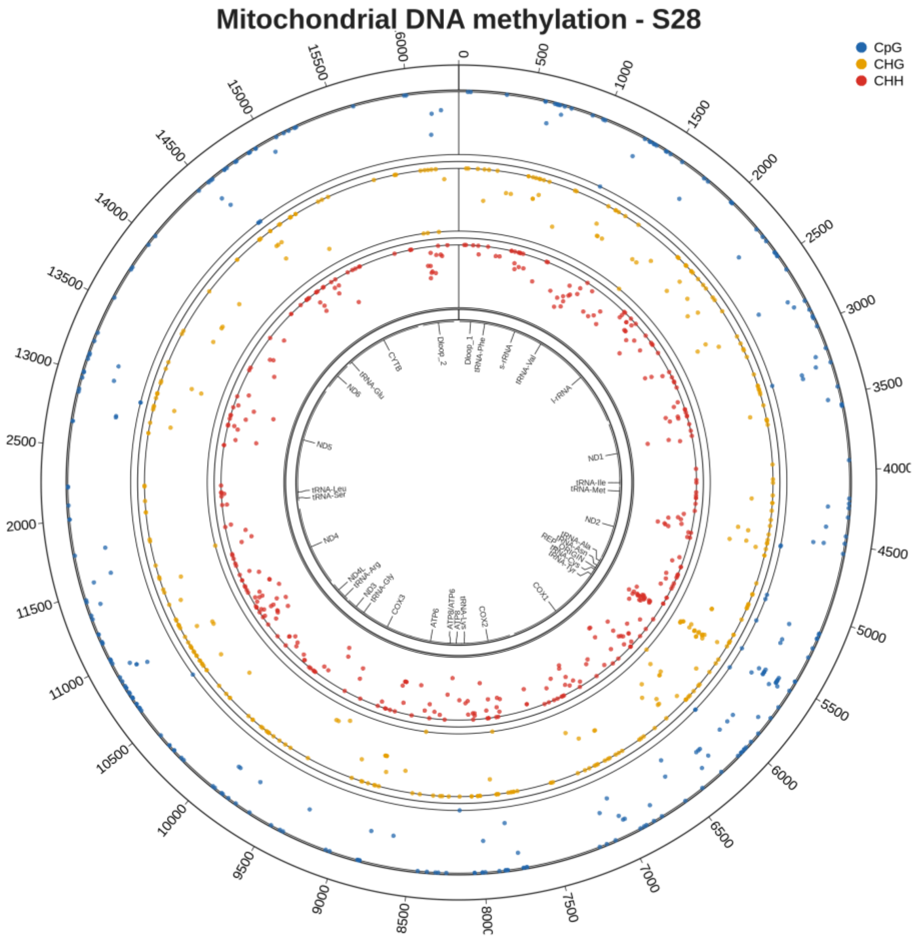
Mitochondrial DNA methylation - S28
CpG
CHG
CHH

Blood

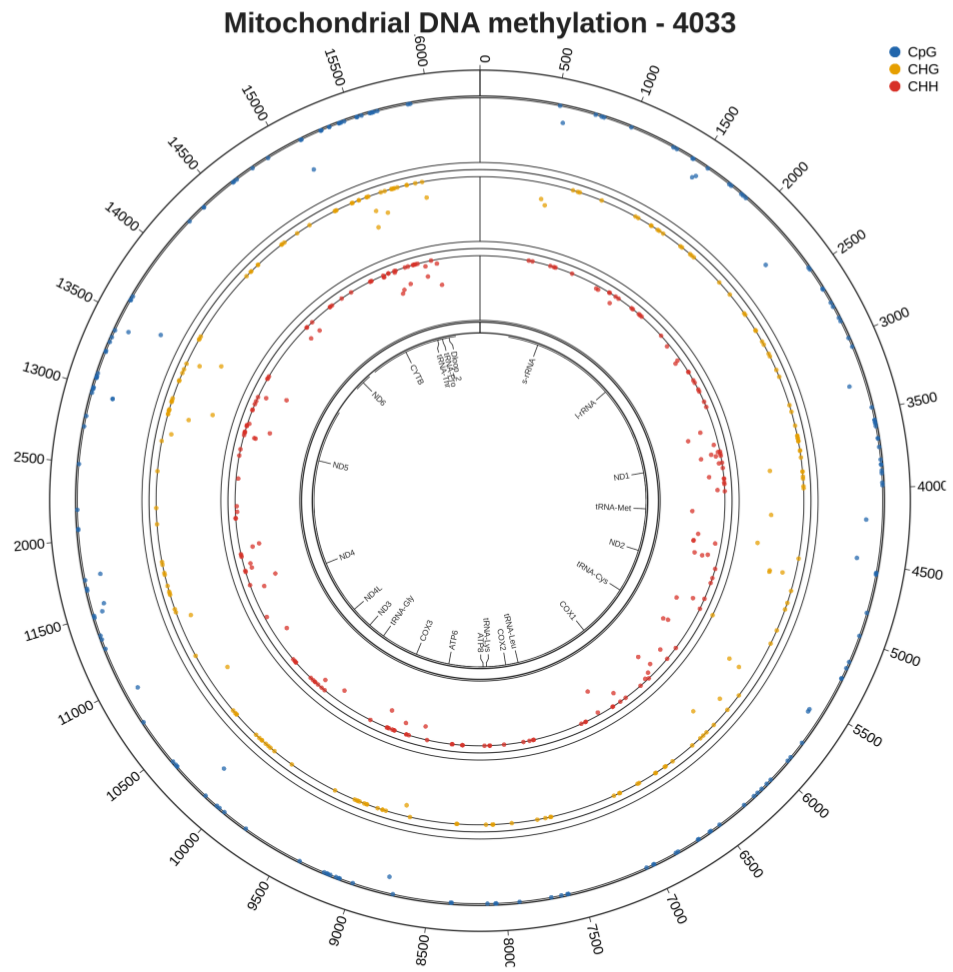
Mitochondrial DNA methylation - 4033
CpG
CHG
CHH
s-rRNA
l-rRNA
ND1
tRNA-Met
ND2
tRNA-Cys
COX1
tRNA-Leu
COX2
ATP8
ATP6
COX3
tRNA-Gly
ND3
ND4L
ND4
ND5
ND6
CYTB
